# The FIP 1.0 Data Set: Highly Resolved Annotated Image Time Series of 4,000 Wheat Plots Grown in Six Years

**DOI:** 10.1101/2024.10.04.616624

**Authors:** Lukas Roth, Mike Boss, Norbert Kirchgessner, Helge Aasen, Brenda Patricia Aguirre-Cuellar, Price Pius Atuah Akiina, Jonas Anderegg, Joaquin Gajardo Castillo, Xiaoran Chen, Simon Corrado, Krzysztof Cybulski, Beat Keller, Stefan Göbel Kortstee, Lukas Kronenberg, Frank Liebisch, Paraskevi Nousi, Corina Oppliger, Gregor Perich, Johannes Pfeifer, Kang Yu, Nicola Storni, Flavian Tschurr, Simon Treier, Michele Volpi, Hansueli Zellweger, Olivia Zumsteg, Andreas Hund, Achim Walter

**Affiliations:** ETH Zürich, Institute of Agricultural Sciences, Zürich, Switzerland; ETH Zürich and EPFL, Swiss Data Science Center, Switzerland; Agroscope, Earth Observation of Agroecosystems Team, Zürich, Switzerland; Universidad Nacional de Colombia, Facultad de Ciencias Agropecuarias, Bogotá, Colombia; University of Wyoming, Department of Plant Sciences, Laramie, Wyoming, USA; ETH Zurich, Plant Pathology Group, Zürich, Switzerland; ETH Zurich, Seminar for Statistics, Zürich, Switzerland; Sítio Florosa, São Paulo, Brazil; The John Innes Centre, Crop Genetics, Norwich, United Kingdom; Agroscope, Water Protection and Substance Flows Team, Zürich, Switzerland; Federal Office for Agriculture and Food, Bonn, Germany; Technical University of Munich, Precision Agriculture Lab, Freising, Germany; Departement für Inneres und Volkswirtschaft (DIV), Landwirtschaftsamt, Thurgau, Switzerland

**Keywords:** Winter wheat, High-throughput phenotyping, Field phenotyping platform, Yield, Protein content, Image time series, Deep learning data set

## Abstract

**Background:** Understanding genotype-environment interactions of plants is crucial for crop improvement, yet limited by the scarcity of quality phenotyping data. This data note presents the Field Phenotyping Platform 1.0 data set, a comprehensive resource for winter wheat research that combines imaging, trait, environmental, and genetic data.

**Findings:** We provide time series data for more than 4,000 wheat plots, including aligned high-resolution image sequences totaling more than 153,000 aligned images across six years. Measurement data for eight key wheat traits is included, namely canopy cover values, plant heights, wheat head counts, senescence ratings, heading date, final plant height, grain yield, and protein content. Genetic marker information and environmental data complement the time series. Data quality is demonstrated through heritability analyses and genomic prediction models, achieving accuracies aligned with previous research.

**Conclusions:** This extensive data set offers opportunities for advancing crop modeling and phenotyping techniques, enabling researchers to develop novel approaches for understanding genotype-environment interactions, analyzing growth dynamics, and predicting crop performance. By making this resource publicly available, we aim to accelerate research in climate-adaptive agriculture and foster collaboration between plant science and machine learning communities.

## Data Description

### Aim

Winter wheat provides a crucial share of calories for human nutrition, with global demand steadily increasing [1]. However, crop production faces challenges due to limited resources like water, agrochemicals, and land [2]. Climate change further threatens crop yields, necessitating responsible and efficient resource use [3].

Crop yields are substantially driven by complex interactions between plant genetics and environmental factors. For instance, genes involved in fruit formation interact with temperatures at flowering, influencing growth and yield potential [4]. Limited phenotyping data is seen as the major reason for the incomplete understanding of such genotype-environment interactions [5].

High-throughput field phenotyping (HTFP) was developed to address this data gap [6]. Imaging HTFP platforms allow researchers to monitor crop canopy development over time, generating dense time series data of plant growth. There are many approaches to process such data ranging from extracting traits at critical time points to modeling growth dynamics and finally using end-to-end methods that directly analyze image time series.

This data set aims to provide a comprehensive foundation for these diverse approaches. Our goal is to foster collaboration between plant physiology, biometrics, and computer vision research, ultimately improving the ability to predict genotype-environment interactions for current and future climates.

### Context

The Field Phenotyping Platform (FIP) at ETH was established in 2015 to collect image time series of crops growing under realistic field conditions. The FIP’s cable carrying system is capable of carrying a 90 kg sensor head [7]. The original sensor head, hereafter referred to as the FIP 1.0 head, was equipped with a red, green, and blue (RGB) camera and a Terrestrial Laser Scanner (TLS), among other sensors. Wheat field experiments were observed using FIP 1.0 over an eight-year period from 2015 to 2022, yielding six years of data collection, with 2015 and 2020 excluded due to incomplete measuring seasons (Figure 1). RGB images of all experimental units (so-called ‘plots’) were collected up to three times a week, and plant heights were measured simultaneously using either the TLS (2016, 2017) [8, 9] or drones (2018–2022) [10, 11, 12], two methods of height measurement that have been in good accordance with one another (*R*^2^: 0.99 [12]). In 2023, the FIP 1.0 sensor head was replaced with a new, multi-view RGB sensor head. The described data set includes all RGB and height data collected in winter wheat experiments up to this replacement.

**Figure 1.**
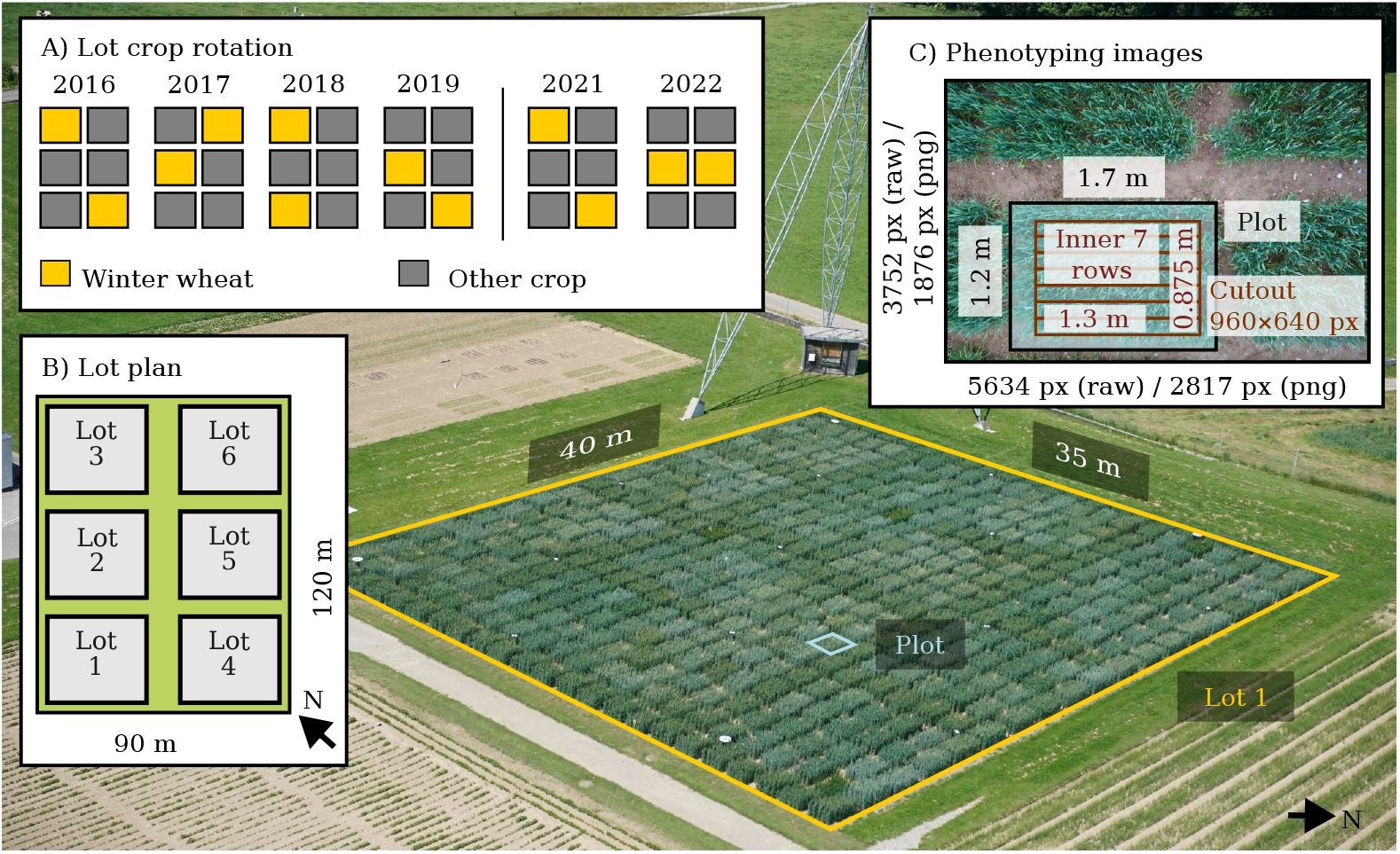
The data set source: 12 wheat lots in 6 years (2016–2022) with >350 wheat plots each, resulting in >4,000 plots from which >160,000 images were taken. The background image shows the Field Phenotyping Platform (FIP) lot 1 with wheat plots. All wheat lots were integrated in a regular crop rotation with other crops (A) according to a permanent lot plan (B). Images taken with the FIP show one complete plot each, the positions of the complete plot and the inner 7 rows are annotated (C).

The area of approximately one hectare that the FIP can monitor is divided into six smaller parts (so-called ‘lots’) that are integrated into a crop rotation. The two FIP lots dedicated to winter wheat provide space for ∼350 genotypes, replicated once per lot. For the first three years (2016–2018), the GABI-WHEAT [13] panel was grown as the genotype set. From 2019–2022, a subset of the GABI-WHEAT panel was grown in addition to other genotypes (Figure 2, green bars). The GABI-WHEAT panel consists of registered genotypes from different climatic regions of Europe [14, 13]. Genetic marker data and Multi-Environment Trial (MET) data from eight year-locations for GABI-WHEAT are publicly available.

**Figure 2.**
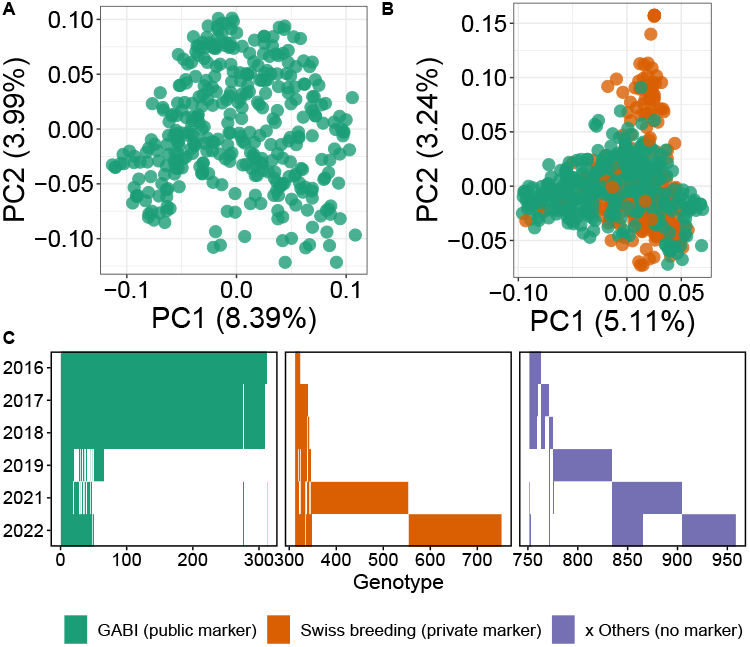
The examined genotype sets with their genetic relatedness visualized in SNP-marker based principal component analysis based on public GABI-WHEAT markers (A) and public and private markers combined (B), and year of cultivation of genotypes (C).

The GABI-WHEAT panel was largely superseded by the Swiss breeding set in 2021 (Figure 2, orange bars). This new set primarily consists of eighth-generation (F8) breeding genotypes. Private genetic marker data are available for the Swiss breeding set. These can be shared directly through the breeders as part of a collaboration. The remaining genotypes, linked to specific projects such as Innovation in Variety Testing (INVITE), were present throughout all years but were generally only grown in a single year each (Figure 2, purple bars). These genotypes currently lack available marker data.

In summary, by default the data set contains public genotypic data from ∼300 genotypes over six years, allowing all the uses described in this data note. Through collaborations, this set can be expanded to ∼800 genotypes.

Regular measurements with the FIP 1.0 head were accompanied by reference measurement campaigns as part of several projects. The heading date and senescence ratings were performed to investigate the relationships of senescence dynamics and diseases [15, 16, 17]. Yield measurements taken on the FIP field were combined with data from other locations to train phenomic prediction models [18]. The plant height measurements served as a basis to quantify the temperature response of wheat genotypes in the stem elongation phase [8, 9, 12]. The extracted plant height values demonstrated their usefulness in improving trait extraction methods from longitudinal data [19, 20, 21].

The images collected allowed to quantify canopy cover values [22] and examine their relationship to frost damage events [23] using Convolutional Neural Networks (CNNs). Using a combination of drone data and the high-resolution images the rows in the individual plots were identified [11]. In a small subset (375 images), the wheat heads were annotated and the data was integrated into the public global wheat head detection data set [24]. The image-based canopy cover values served as a test data set to evaluate the cultivar-specific extensions of the thermal time concept [25].

The culmination of these efforts has resulted in a unique, multi-dimensional data set. Dense image time series of diverse wheat genotypes are integrated with trait measurements, genetic markers, and environmental data. The data set has been designed to align with FAIR principles [26]:

- **Findable**: This publication and the Hugging Face data set card (https://doi.org/10.57967/hf/3191) provide detailed metadata and a comprehensive description of the data set’s contents, making it discoverable to researchers.
- **Accessible**: The data is hosted on the Research Collection of ETH Zurich (https://doi.org/20.500.11850/697773), a reliable and openly accessible data storage.
- **Interoperable**: The use of the open-source Hugging Face datasets [27] package makes it easy to use and export to different formats. The data is fully MIAPPE v1.1 [28] conform. Given the shared genotypes the data set can be used to enhance the data by Gogna et al. [13] by 6 environments, to a total of 14 environments. The data set expands on existing sub-sets of the data already released that can be used as baseline approaches such as [8, 9, 19, 20, 21, 12, 25, 15, 16, 17].
- **Reusable**: The data is released under the CC0 1.0 Univer-sal license (https://creativecommons.org/publicdomain/zero/1.0/), a permissive license that allows for further use of the data.

## Materials and Methods

### Experimental Field Designs and Genotypes

All experiments were performed at the ETH research station of plant sciences in Lindau Eschikon, Switzerland (47.449 N, 8.682 E, 556 m a.s.l.). The soil characteristics were determined in 2015 (Eric Schweizer AG, Thun, Switzerland). The soil type is eutric cambisol consisting of 21% clay and 21% silt with an organic matter content of 3.5% and pH 6.7. A crop rotation was implemented during and before the start of wheat experiments beginning with a year of soybean (*Glycine max* (L.) Merr.), then a year of buckwheat (*Fagopyrum esculentum* Moench) and finally wheat (*Triticum aestivum* L.). After preliminary crops were harvested, the soil was plowed and harrowed before wheat was drill-sown. The wheat was sown in 9 rows per plot with a row length of ∼1.7 m, a row distance of 0.125 m and a sowing density of 370–400 plants m^-2^.

A few days after sowing, herbicide (Herold SC, Bayer AG, Leverkusen, Germany) was applied to ensure weed-free plots. Several fungicides and insecticides were applied in spring to ensure healthy plants. The fertilizer was split into three doses (∼ 1:3:1), one at tillering stage, one at start of stem elongation, and one after heading. Approximately 140 kg N, 90 kg P_2_O_5_, and 100 kg K_2_O per ha were applied, depending on site-specific soil analysis. No irrigation was applied. The sowing and harvest dates for each year are provided in Table 1.

**Table 1.** Sowing and harvest dates.

| Year | Sowing Date | Harvest Date |
| --- | --- | --- |
| 2016 | 2015-10-13 | 2016-07-27 |
| 2017 | 2016-11-01 | 2017-07-19 |
| 2018 | 2017-11-02 | 2018-07-14 |
| 2019 | 2018-10-17 | 2019-07-23 |
| 2021 | 2020-10-21 | 2021-07-29 |
| 2022 | 2021-11-25 | 2022-07-19 |

For 2016–2018, a GABI-WHEAT panel subset (consisting of ∼300 European winter wheat cultivars from the GABI-WHEAT panel [14, 29]) was complemented by 35–52 Swiss winter wheat varieties of commercial importance. For 2019, a small subset of the GABIWHEAT panel (54 genotypes) was grown alongside genotypes from other experiments (e.g., Swiss variety testing). In 2019, these other genotypes were grown on larger plots with a row length of 5.5 m while GABI-WHEAT genotypes were grown on the same size plots as in the other years. In 2021 and 2022, sets of F8 genotypes from the Swiss breeding program of Agroscope (Nyon, Switzerland) were grown. For an overview of genotype overlaps between years, see Figure 2.

For all years, an experimental design following the principles of good practice—replication, randomization, and blocking [30]— was chosen. The described panels of, on average, 350 genotypes per year were replicated once, each replication was randomized and grown on a different lot in the FIP area (Figure 1). Each replication was augmented with checks in a 3×3 block arrangement. For further details on the experimental design, see [8, 9, 12].

### Image Data

The FIP system is divided into two independent parts, (1) the carrier system built and maintained by Spidercam (Spidercam GmbH, Feistritz im Rosental, Austria), and (2) the custom-built FIP 1.0 sensor head and control software [7].

#### Carrier: The Spidercam Cable-Suspended System

The carrier system, detailed in Kirchgessner et al. [7], uses four corner-mounted poles with pulleys. Cables connect these pulleys to winches, enabling 3-D movement of the FIP 1.0 sensor head. A working distance of 2–3 m from the canopy was maintained during measurements.

#### Sensors: The FIP 1.0 Imaging Head

The FIP 1.0 sensor head carried, amongst other sensors, a 21 MP full frame DSLR camera (EOS 5D Mark II, 35 mm lens (Canon Inc., Tokyo, Japan) [7]. The camera was triggered automatically via a custom MATLAB script (The Mathworks [31]). The images were mostly captured using auto white balance, auto exposure, an ISO of 100, an exposure time of 1/250 second (4 ms), and zero exposure bias value. The ground sampling distance is approximately 0.55 mm.

#### Image Registration

The positions of the captured images of the plots change throughout the season due to inaccuracies in the camera carrier system and intentional height adjustments due to plant growth. To allow for consistent image feature extraction, the time series needed to be aligned using an image registration pipeline. Image registration transforms the images so that points that are at the same physical location in the real world are aligned to the same point in the image planes. The used image registration pipeline consists of two steps: a deep learning-based feature matcher and a transformation estimation step. The feature matcher is used to find features that correspond to the same location between an image pair, which are then used to estimate the transformation between the images. The registration was performed between image pairs instead of the whole sequence directly, commonly referred to as image alignment.

To predict aligned polygons of the inner rows (Figure 1) for an image time series three individual steps were performed. First an initial reference polygon was aligned with a drone orthomosaic or prior data. Then subsequent aligned polygons were found based on this initial polygon or a previously aligned polygon. Finally, a single transformation between alignments and the inner plots was estimated and applied to all aligned polygons. The aligned result of a time series can be seen in Figure 3.

**Figure 3.**
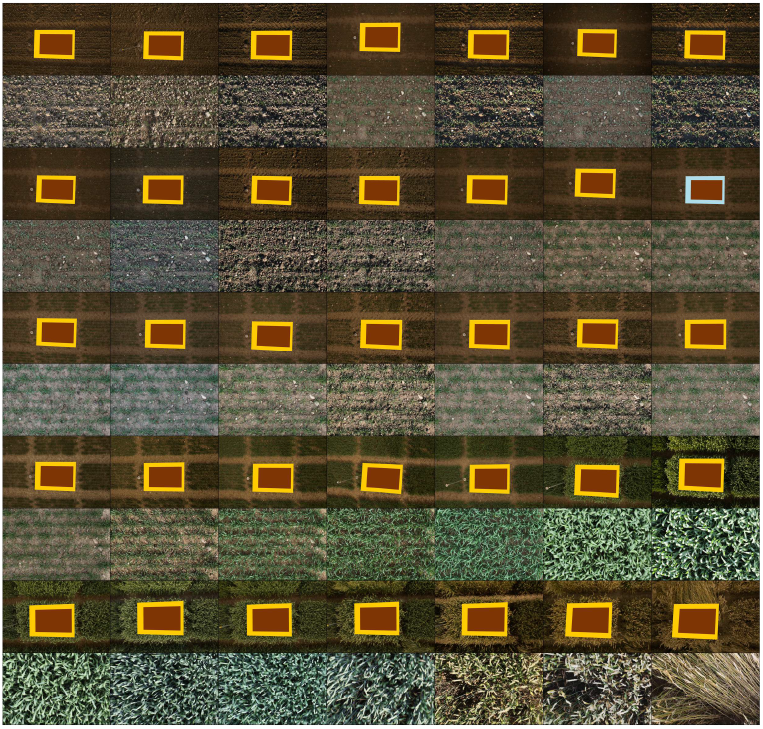
An image time series of a plot over one season with the aligned inner plot cutouts in each second row. The initial alignment polygon, shown in light blue, is based on alignment with a drone orthophoto. All other alignment polygons, shown in orange, are transformed from the initial polygon or another aligned polygon. The inner plot polygons, shown in brown, are based on a single transform from an aligned polygon to the inner rows of the plot. Images are adjusted for visibility.

To find the initial reference polygon, the 3D-world-coordinates of the plot corners were first extracted from drone-based orthomosaics (2018–2022), from other projects [32, 10, 11, 33, 18, 12], or from extrapolations of these plot corners to earlier years (2016– 2017). The 3D plot corners from this initial extraction process were matched to the best-fitting image in the image time series to find their 2D counterparts, which were used as the initial reference polygon.

Using the initial reference image and polygon, the initial image was matched with other images and the resulting transformation used to find the other aligned plot corners. Matching features between different time points in a crop season is challenging due to significant changes in conditions between images, such as varying lighting, plant growth and changing appearance of the plants. For this reason, a modified version of the deep learning LoFTR [34] feature matcher was fine-tuned to focus on consistent soil features, such as stones. Even these consistent objects slowly change their positions throughout the season or rapidly between strong precipitation events. Therefore, the transformations were estimated between image pairs and not the complete time series at a time. The estimated transformations between the image pairs are homographies based on the soil plane.

A prior alignment strategy employing Scale-invariant feature transform (SIFT) based feature matching required manual alignment of about 10,000 image pairs to fill gaps where the matcher failed. While suitable for smaller scale projects, this approach was deemed unviable for the complete data set. These manual alignments, created by choosing four point-pairs to find suitable homographies, were used to fine-tune the LoFTR feature matcher. The homographies correspond to the transformations between the common soil plane of the image pairs. This simplification was used to fine-tune the model to find matches solely in the soil plane, as features in the canopy are at incorrect positions due to not being on the homography plane. Since the canopy moves, sometimes significantly, between time-points, this reduces the number of incorrect matches that the feature matcher produces. The model was fine-tuned for 20 epochs with randomly cropped, rotated and distorted image pairs. These augmentations were applied to each image individually, adjusting the homographies accordingly. The augmentations were chosen such that the image pairs were always partially overlapping.

To verify the correctness of the matches and their homographies, several checks were employed. Basic checks, such as ensuring enough inliers, constraining the ratios of the side lengths and angles of the polygon spanned by the plot corners to their 3D counterparts, and other image-based checks were applied with high thresholds. The essential matrix was calculated, and the warped plot corners were compared to their closest point on the epipolar line. The essential matrix was used to triangulate the 3D points of the plot corners, and they were compared to the original 3D-world-coordinates by checking the ratios and angles of the polygons spanned by the plot corners. These ratios were used to scale the predicted relative camera movement and compare it individually to the maximum possible movements in each axis. Finally, these predictions and checks were done twice with the image pair swapped and compared to each other. This check filtered cases where LoFTR predicted structured noise that corresponded to the identity homography between the image pairs. To filter outliers these steps were iteratively repeated, and the best prediction based on the checks was chosen.

When an image could not be matched to the initial image, it was iteratively matched with the closest found alignment, based on the number of transformations and then the date differences. Matches were used to find homographies using OpenCV [35], which were then used to warp the plot corners (Figure 1, ‘Plot’).

The complete process, except the initial 3D-world-coordinates extraction, was iteratively repeated by going from very tight to looser restrictions in the checks. The fine-tuned LoFTR model was bootstrapped, that is, further trained at each step by adding the trusted subset of its predictions that passed the checks in the previous step to the training set.

Finally after predicting all plot corners, a relative inner plot polygon was extracted to mitigate border effects and further correct the alignment. The seven inner rows of plants were detected based on segmented images showing plant and soil pixels. Then, plots were further rectified by rotating them step-wise (−1.5° to 1.5° in steps of 0.2°) to maximize the distance between the minimum and maximum numbers of plant pixels in image columns [25]. Inner plots were filtered if they did not contain the complete plot.

In total, the alignment and inner plot detection was successful for more than 95% of all images. The number of successfully aligned images per year can be seen in Table 2. Both the 160,772 original images and their corresponding 153,022 inner plot cutouts are made available as part of this data set.

**Table 2.** Data set sizes for original images and aligned images per year. For each category, the number of images and minimum / maximum / mean time-series length are given.

| Year | Plots | Original images |  |  |  | Aligned images |  |  |  |
| --- | --- | --- | --- | --- | --- | --- | --- | --- | --- |
|  |  | # Images | Min | Max | Mean | # Images | Min | Max | Mean |
| 2016 | 710 | 26237 | 35 | 39 | 36.95 | 25566 | 26 | 39 | 36.01 |
| 2017 | 756 | 19717 | 23 | 30 | 26.08 | 14534 | 9 | 29 | 19.22 |
| 2018 | 756 | 21642 | 26 | 31 | 28.63 | 21266 | 12 | 31 | 28.13 |
| 2019 | 252 | 7451 | 28 | 34 | 29.57 | 6445 | 20 | 34 | 25.57 |
| 2021 | 792 | 48451 | 58 | 63 | 61.17 | 48285 | 58 | 63 | 60.97 |
| 2022 | 792 | 37274 | 44 | 51 | 47.06 | 36926 | 44 | 51 | 46.62 |
| Total | 4058 | 160772 | 23 | 63 | 39.62 | 153022 | 9 | 63 | 37.71 |

#### Image Preprocessing

The raw images were converted to PNG format, an accessible and lossless image format, using rawpy [36] with minimal post-processing, reducing resolution to half size, employing a linear demosaicing algorithm, and disabling auto adjustments in order to preserve the original sensor data with minimal artifacts or alterations. The inner plots were directly cut out of the pre-processed image by combining the plot and its relative inner plot transformation.

### Reference Measurements

As part of several projects [8, 9, 15, 16, 18, 12], reference measurements such as grain yield and growth stage ratings were taken. Those traits can be divided into low-level traits (time series of traits that develop over time), intermediate traits (traits extracted from low-level traits that describe the growth dynamics over time [20, 19]) and target traits (observations that are usually targeted in breeding and agriculture, e.g., yield) (Table 7). Trait measurements may have been made at times different from the image captures. All measurements are scalars that correspond to a single measurement at a given time point during the season.

#### Low-level Traits

Plant height measurements were performed with a TLS (Focus 3D S 120, 905 nm laser, Faro Technologies Inc., Lake Mary, USA) for 2016 and 2017 on the same date as image acquisition [8]. From 2019– 2022, drone RGB images [10, 11] were used to extract plant height estimations with Structure from Motion (SfM) [37]. FIP image collection dates and drone campaign dates were typically within days of each other but did not necessarily overlap. The accuracy of TLS and drone measurements to approximate manual plant height measurements were demonstrated in the respective publications (TLS [8]: *R*^2^: 0.99, drone [37]: *R*^2^: 0.96). The two methods are in good accordance with one another (*R*^2^: 0.99, intercept: 0.057 m, slope: 1.0 [12]).

From images, canopy cover was extracted using a deep learning model described in Zenkl et al. [22], a model that reached a pixel accuracy of 0.945 on a FIP test set. The percentage of soil covered by plant parts was determined for the seven-row inner plot (Figure 1) as described in Tschurr et al. [38]. On the same seven-row inner plot, wheat head count estimations were determined for all dates from May to end of season. As the wheat head detection method, the winning model of the global wheat head challenge [24] (https://github.com/ksnxr/GWC_solution) was used, a model that achieved an average domain accuracy of 0.7 on a test set that included FIP images.

Senescence was assessed visually in 2016, 2017 and 2018 from approximately 20 days after flowering to full senescence for the central plot area canopy, following guidelines provided by Pask et al. [39]. Plot senescence was scored according to Anderegg et al. [16] based on the portion of green leaf area on a scale from 0 to 10, equivalent to 0 to 100 %. To avoid bias, manual ratings were performed by the same expert in all three years.

The number of total measurements, as well as the minimum, maximum, and mean number of measurements per year for each low-level trait, can be seen in Table 3.

**Table 3.** Time-series data set sizes for low-level traits. For each low-level trait the number of measurements and minimum / maximum / mean time-series length are given.

| Year | Plots | Canopy Cover |  |  |  | Plant Height |  |  |  | Wheat head count |  |  |  | Senescence rating |  |  |  |
| --- | --- | --- | --- | --- | --- | --- | --- | --- | --- | --- | --- | --- | --- | --- | --- | --- | --- |
|  |  | # | Min | Max | Mean | # | Min | Max | Mean | # | Min | Max | Mean | # | Min | Max | Mean |
| 2016 | 710 | 710 | 27 | 39 | 36.00 | 710 | 22 | 22 | 22.00 | 710 | 1 | 11 | 9.51 | 703 | 0 | 9 | 8.78 |
| 2017 | 756 | 756 | 9 | 24 | 18.52 | 756 | 21 | 21 | 21.00 | 756 | 1 | 18 | 9.77 | 756 | 1 | 10 | 9.77 |
| 2018 | 756 | 756 | 12 | 27 | 21.50 | 756 | 36 | 43 | 39.72 | 755 | 0 | 18 | 10.35 | 756 | 12 | 12 | 12.00 |
| 2019 | 252 | 252 | 8 | 25 | 21.89 | 252 | 38 | 46 | 44.55 | 228 | 0 | 10 | 3.57 | — | — | — | — |
| 2021 | 792 | 792 | 55 | 61 | 59.21 | 792 | 45 | 48 | 46.56 | 792 | 30 | 36 | 33.96 | — | — | — | — |
| 2022 | 792 | 792 | 31 | 41 | 38.46 | 792 | 42 | 45 | 42.53 | 792 | 12 | 24 | 21.55 | — | — | — | — |
| Total | 4058 | 4058 | 8 | 61 | 34.18 | 4058 | 21 | 48 | 35.13 | 4057 | 0 | 36 | 16.47 | 2215 | 0 | 12 | 5.59 |

#### Intermediate Traits

Heading date was visually assessed as the date when 50% of the spikes were fully emerged from the flag leaf sheath [17] (BBCH 59, [40]). To ensure consistent ratings over time, heading date ratings were started at approximately BBCH 55 and continue to BBCH 61, with two to three rating events per week.

Final height was extracted from TLS or SfM plant height measurements using the Quarter of Maximum Elongation Rate (QMER) method described in Roth et al. [20], which has a reported accuracy of close to 1.0 on simulated data. The number of measurements per year for intermediate and target traits can be seen in Table 4.

**Table 4.**
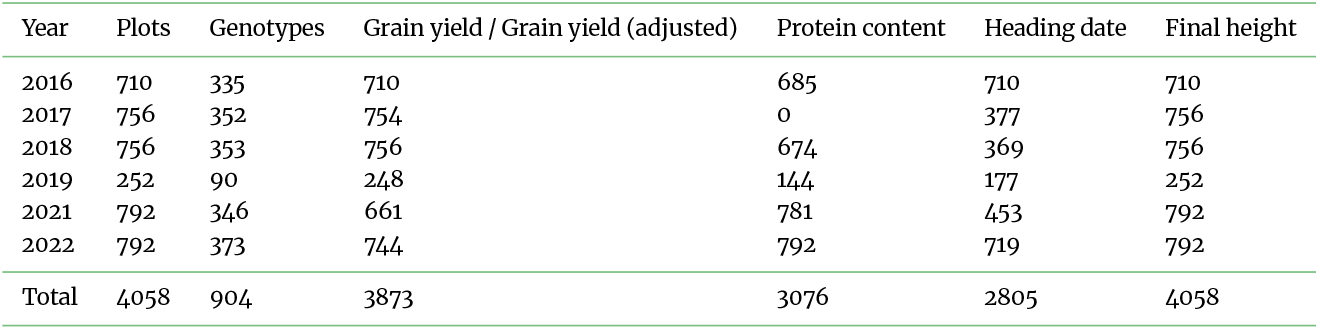
Data set sizes and key characteristics for genotypes as well as intermediate and target traits. Numbers describe absolute sizes, i.e., how many plots and genotypes were examined in a specific year, and how many of this plots/time series are annotated with target traits grain yield, grain protein content, heading date, and final height.

| Year | Plots | Genotypes | Grain yield / Grain yield (adjusted) | Protein content | Heading date | Final height |
| --- | --- | --- | --- | --- | --- | --- |
| 2016 | 710 | 335 | 710 | 685 | 710 | 710 |
| 2017 | 756 | 352 | 754 | 0 | 377 | 756 |
| 2018 | 756 | 353 | 756 | 674 | 369 | 756 |
| 2019 | 252 | 90 | 248 | 144 | 177 | 252 |
| 2021 | 792 | 346 | 661 | 781 | 453 | 792 |
| 2022 | 792 | 373 | 744 | 792 | 719 | 792 |
| Total | 4058 | 904 | 3873 | 3076 | 2805 | 4058 |

**Table 5.** Heritabilities (*H*^2^ ) of intermediate and target traits for all years.

| Trait | Heritability ($H^2$) | | | | | |
| --- | --- | --- | --- | --- | --- | --- |
|  | 2016 | 2017 | 2018 | 2019 | 2021 | 2022 |
| Grain yield | 0.58 | 0.36 | 0.33 | 0.79 | 0.41 | 0.48 |
| Grain yield (adjusted) | (s) | (s) | (s) | 0.83 | (s) | (s) |
| Protein content | 0.82 | - | 0.81 | 0.80 | 0.55 | 0.86 |
| Heading date | 0.96 | N/A | 0.90 | 0.89 | 0.81 | 0.93 |
| Final height | 0.97 | 0.98 | 0.97 | 0.93 | 0.96 | 0.97 |

N/A: Unreplicated measurement, heritability not available
–: No measurements in this year
Grain yield (adjusted): Plot size adjusted yield ( $\text{yield}_{5\text{m}} = \text{yield}_{1\text{m}} / 1.50$ )
(s): All plot same size, see results for “Grain yield”

#### Target Traits

Yield for small plots in all years was estimated based on two rows of nine rows (row seven and eight). Ears within these two rows were hand-harvested, dried for at least 24 hours at 30 °C, and threshed using a stand thresher (Saatmeister Allesdrescher K35; Saatzucht Baumann, Germany). Yield for large plots in 2019 was determined with a combine harvester (Nursery-master Elite; Wintersteiger, Ried im Innkreis, Austria). Weight was determined using a scale, and water content with a Wile 55 moisture meter (Farmcomp Oy; FIN04360 Tuusula, Finland). Grain yield was mathematically normalized to 14 % water content. Grain protein content was determined using near-infrared transmission spectroscopy (InfratecTM 1241 Grain Analyzer; Foss, DK-3400 Hilleroed, Denmark).

In 2019, plot sizes varied between experiments, which influences yield measurements. While large plot sizes deliver absolute estimates comparable with METs, in our experience, small plot sizes tend to overestimate yield per area. To compensate for this effect, adjusted genotype means (see next Section) of twelve common genotypes between the experiments with large and small plots were used to calculate a conversion factor. A linear regression with intercept zero estimated a conversion factor of 1.50 from large to small plots (*R*^2^ of 0.47). This conversion factor was used to transform all yield measurement estimates of small plots to the range of large plots, resulting in a new trait ‘Grain yield (adjusted)’.

#### Adjusted Genotype Means within Year Calculation

All intermediate and target traits were processed to adjusted genotype means (Best Linear Unbiased Estimate (BLUE)) using a linear mixed model in SpATS,

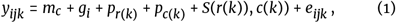

where *y*_*ijk*_ is the measured trait value for the *i*th genotype in the *j*th year for plot *k* in row *r*(*k*) and column *c*(*k*). *m*_*c*_ is a fixed effect marking check varieties (*m*_*c*_ ∈ [0, 1]), *g*_*i*_ is a fixed genotype effect, *p*_*r*(*k*)_ and *p*_*c*(*k*)_ are random spatial row and column effects, and *e*_*ijk*_ a spatially independent residual. *S*(*r*(*k*)), *c*(*k*)) is a spatial smooth surface in row and column direction as defined in [41].

#### Environmental Covariates

Air temperature, relative humidity, short wavelength solar irradiance, and soil temperature were measured at a local weather station in proximity to the experimental field above or below a grass strip [25]. Air temperature and relative humidity were measured 2 m and 0.1 m above ground, short wavelength solar irradiance 2 m above ground, and soil temperature 0.05 m below ground, respectively. Precipitation data were taken from a close-by Agrometeo weather station at Strickhof (<800 m, https://www.agrometeo.ch). Measurement gaps caused by technical issues (e.g., sensor failure) were filled with data from the Strickhof station if available and with data from a Meteoswiss station at the Zurich Airport (9.5 km, https://gate.meteoswiss.ch/idaweb/) otherwise. Overall, more than 97% of the data after gap filling originated from the local weather station or Strickhof weather station.

### Genetic Marker Data

Genetic marker data (90k SNP array) for the GABI-WHEAT panel are publicly available [29]. The private marker data were created using a 25k SNP array. Two sets were compiled: A pure GABI-WHEAT marker data set, and an extended marker set consisting of overlapping marker from both public and private markers. Marker locations on the reference genome IWGSC RefSeq v1.0 were collected with InterMine [42] from https://urgi.versailles.inrae.fr and complemented with locations previously mapped with blastn from an earlier project [12].

SNPs in both sets were first filtered for minor allele frequency (5%) and missing values (5%). Genotypes in both sets were then tested for missing marker data with a missing rate of 1% (all passed, no genotype had to be removed). Finally, missing marker data (0.44% for GABI-WHEAT set, 0.54% for extended set) were imputed based on a k-nearest-neighbour implementation in R (scrime [43]) per chromosome. Markers were sorted by chromosome and location for later use in local connectivity models such as CNNs. Markers with ambiguous positions were left in the set but marked accordingly.

The resulting GABI-WHEAT genetic marker set includes 372 genotypes and 18’846 markers. The extended set includes 824 genotypes and 11’943 markers. Based on the marker data, kinship matrices [44] were calculated according to Yang [45].

### Compilation as Data Set

#### Train/Test Split According to Genetic Relatedness

The most interesting application of trait prediction approaches is to predict the performance of unseen genotypes in unseen years [46, 47]. The accuracy of such predictions typically depends on the relatedness of genotypes, usually assessed through cross-validation. However, cross-validation is computationally expensive for complex deep learning models. To address this issue, we propose an alternative approach that balances the train/test set using genetic relatedness [48]. This method theoretically yields performance close to the average of all cross-validation runs.

To implement this approach, we used the R package STPGA [49] to determine a test set complementing the training set with the algorithm ‘D_opt_’ as suggested in [50]. The training set was further split into training and validation sets using the same method. We define four test sets to allow for specific evaluation of methods depending on their use. The test sets differ based on if their genotypes and environments occur in the train set (denoted as seen):

i. Test (P): Unseen plots with seen genotypes and seen environments (all years except 2019)
ii. Test (G): Unseen genotypes with seen environments
iii. Test (E): Unseen environment (2019) with seen genotypes
iv. Test (G and E): Unseen genotypes and unseen environment (2019)

To validate our balanced splitting approach, we also performed five-fold cross-validations, allowing for a quality check of the balanced splits. The splits were calculated separately for the pure GABIWHEAT set and for the extended set. The extended set additionally includes F8 generation genotypes with private marker data. For the GABI-WHEAT set, this resulted in 262 genotypes in the training set, 24 in the validation set, and 30 in Test (G and E) test set (Figure 5). For the extended set, this resulted in 747 genotypes in the training set, 30 in the validation set, and 30 in Test (G and E) test set.

#### Preparation as Hugging Face Data Set

All data was aggregated into a single table with a row for each plot containing the image sequence, the aligned image sequence, traits, environmental data, marker data, and additional metadata. The table was split using the aforementioned splits and converted to a single Hugging Face data sets DatasetDict using the schema shown in Table 8. The table and DatasetDict contain None values for completely missing entries. Missing data in sequences are absent.

**Table 6.** Variance decomposition results of intermediate and target traits for all years.

| Trait | Component | Percentage of total variance |  |  |  |  |  |
| --- | --- | --- | --- | --- | --- | --- | --- |
|  |  | 2016 | 2017 | 2018 | 2019 | 2021 | 2022 |
| Grain yield | Genotype | 8% | 11% | 1% | 1% | 18% | 3% |
|  | Spatial | 81% | 52% | 94% | 98% | 41% | 91% |
|  | Residual | 11% | 37% | 5% | 1% | 41% | 6% |
| Grain yield (adjusted) | Genotype | (s) | (s) | (s) | 28% | (s) | (s) |
|  | Spatial | (s) | (s) | (s) | 59% | (s) | (s) |
|  | Residual | (s) | (s) | (s) | 13% | (s) | (s) |
| Protein content | Genotype | 1% | - | 2% | 11% | 7% | 6% |
|  | Spatial | 98% | - | 97% | 85% | 83% | 92% |
|  | Residual | 0% | - | 1% | 4% | 11% | 2% |
| Heading date | Genotype | 61% | N/A | 58% | 57% | 77% | 35% |
|  | Spatial | 34% | N/A | 36% | 29% | 2% | 61% |
|  | Residual | 4% | N/A | 6% | 14% | 20% | 4% |
| Final height | Genotype | 60% | 87% | 5% | 19% | 6% | 12% |
|  | Spatial | 36% | 9% | 95% | 77% | 94% | 88% |
|  | Residual | 3% | 3% | 0% | 3% | 0% | 1% |

N/A: Unreplicated measurement, variance components not available
-: No measurements in this year
Grain yield (adjusted): Plot size adjusted yield ( $\text{yield}_{5\text{ m}} = \text{yield}_{1\text{ m}} / 1.50$ )
(s): All plots same size, see results for “Grain yield”

**Table 7.** Measured traits that were used to annotate image time series. Label refers to the name used in the data set for the trait.

| Trait | Unit | Label | Description |
| --- | --- | --- | --- |
| **Low-level traits** |  |  |  |
| Canopy cover | (0...1) | canopy_cover | Soil coverage by plants from a nadir view |
| Plant height | m | height | Distance top of soil to top of canopy |
| Wheat head count | Count | spike_count | Number of visible wheat spikes |
| Senescence rating | (0...10) | senescence | End-of-season leaf decay ratings |
| **Intermediate traits** |  |  |  |
| Heading | Date | heading | Date when 50% of wheat spikes visible |
| Final height | m | height_final | Plant height at end-of-season |
| **Target traits** |  |  |  |
| Grain yield | t/ha | yield | Total weight of harvested grains |
| Grain yield (adjusted) | t/ha | yield_adjusted | Plot size adjusted yield |
| Protein content | % | protein | Grain protein content |

**Table 8.** A schema of the data set equivalent to the Hugging Face data set schema. Sequence indicates a variable length of values while Array2D indicates a fixed shape of the data.

| Field Name | Type | Field Name | Type |
| --- | --- | --- | --- |
| **identifiers** |  | **height_final_value** | float16 |
| plot_uid | string | **height_final_date** | date32 |
| yearsite_uid | string | **height_final_blue** | float16 |
| crop_type | string | **height_final_heritability** | float16 |
| experiment_number | uint8 | **height_final_trait_id** | uint8 |
| plot_number | int16 | **height_final_trait_name** | string |
| **location** |  | **height_final_method_id** | uint8 |
| range | uint8 | **height_final_method_name** | string |
| row | uint8 | **height_final_si_unit** | string |
| lot | uint8 | **height_final_responsible** | string |
| latitude | float64 | **target traits** |  |
| longitude | float64 | **yield_value** | float16 |
| spatial_check | int32 | **yield_date** | date32 |
| **dates** |  | **yield_blue** | float16 |
| sowing_date | date32 | **yield_heritability** | float16 |
| harvest_date | date32 | **yield_trait_id** | uint8 |
| harvest_year | uint16 | **yield_trait_name** | string |
| **images** |  | **yield_method_id** | uint16 |
| images | Sequence[string] | **yield_method_name** | string |
| image_dates | Sequence[date32] | **yield_si_unit** | string |
| image_times | Sequence[time32[s]] | **yield_responsible** | string |
| **alignments** |  | **yield_adjusted_value** | float16 |
| alignment_plot_soil_polygons | Sequence[Array2D[float16]] | **yield_adjusted_date** | date32 |
| alignment_num_steps | Sequence[uint8] | **yield_adjusted_blue** | float16 |
| alignment_dates | Sequence[date32] | **yield_adjusted_heritability** | float16 |
| alignment_times | Sequence[time32[s]] | **yield_adjusted_trait_id** | uint8 |
| alignment_initial_date | date32 | **yield_adjusted_trait_name** | string |
| alignment_inner_plot_transform | Array2D[float16] | **yield_adjusted_method_id** | uint16 |
| inner_plot_images | Sequence[string] | **yield_adjusted_method_name** | string |
| image_inner_plot_transforms | Sequence[Array2D[float16]] | **yield_adjusted_si_unit** | string |
| **markers** |  | **yield_adjusted_responsible** | string |
| genotype_id | string | **protein_value** | float16 |
| marker_biallelic_codes | Sequence[uint8] | **protein_date** | date32 |
| marker_metadata_strings | Sequence[string] | **protein_blue** | float16 |
| **low-level traits** |  | **protein_heritability** | float16 |
| canopy_cover_values | Sequence[float16] | **protein_trait_id** | uint8 |
| canopy_cover_dates | Sequence[date32] | **protein_trait_name** | string |
| canopy_cover_trait_ids | Sequence[uint8] | **protein_method_id** | uint16 |
| canopy_cover_trait_name | Sequence[string] | **protein_method_name** | string |
| canopy_cover_method_ids | Sequence[uint16] | **protein_si_unit** | string |
| canopy_cover_method_name | Sequence[string] | **protein_responsible** | string |
| canopy_cover_si_unit | string | **environment** |  |
| canopy_cover_responsible | string | **temperature_air_10cm_values** | Sequence[float16] |
| height_values | Sequence[float16] | **temperature_air_10cm_dates** | Sequence[date32] |
| height_dates | Sequence[date32] | **temperature_air_10cm_times** | Sequence[time32[s]] |
| height_trait_ids | Sequence[uint8] | **temperature_air_200cm_values** | Sequence[float16] |
| height_trait_name | Sequence[string] | **temperature_air_200cm_dates** | Sequence[date32] |
| height_method_ids | Sequence[uint16] | **temperature_air_200cm_times** | Sequence[time32[s]] |
| height_method_name | Sequence[string] | **temperature_soil_5cm_values** | Sequence[float16] |
| height_si_unit | string | **temperature_soil_5cm_dates** | Sequence[date32] |
| height_responsible | string | **temperature_soil_5cm_times** | Sequence[time32[s]] |
| spike_count_values | Sequence[float16] | **humidity_air_10cm_values** | Sequence[float16] |
| spike_count_dates | Sequence[date32] | **humidity_air_10cm_dates** | Sequence[date32] |
| spike_count_trait_ids | Sequence[uint8] | **humidity_air_10cm_times** | Sequence[time32[s]] |
| spike_count_trait_name | Sequence[string] | **humidity_air_200cm_values** | Sequence[float16] |
| spike_count_method_ids | Sequence[uint16] | **humidity_air_200cm_dates** | Sequence[date32] |
| spike_count_method_name | Sequence[string] | **humidity_air_200cm_times** | Sequence[time32[s]] |
| spike_count_si_unit | string | **precipitation_200cm_values** | Sequence[float16] |
| spike_count_responsible | string | **precipitation_200cm_dates** | Sequence[date32] |
| senescence_values | Sequence[float16] | **precipitation_200cm_times** | Sequence[time32[s]] |
| senescence_dates | Sequence[date32] | **irradiance_solar_200cm_values** | Sequence[float16] |
| senescence_trait_ids | Sequence[uint8] | **irradiance_solar_200cm_dates** | Sequence[date32] |
| senescence_trait_name | Sequence[string] | **irradiance_solar_200cm_times** | Sequence[time32[s]] |
| senescence_method_ids | Sequence[uint16] |  |  |
| senescence_method_name | Sequence[string] |  |  |
| senescence_si_unit | string |  |  |
| senescence_responsible | string |  |  |
| **intermediate traits** |  |  |  |
| heading_value | float16 |  |  |
| heading_date | date32 |  |  |
| heading_blue | float16 |  |  |
| heading_heritability | float16 |  |  |
| heading_trait_id | uint8 |  |  |
| heading_trait_name | Sequence[string] |  |  |
| heading_method_id | uint16 |  |  |
| heading_method_name | Sequence[string] |  |  |
| heading_si_unit | string |  |  |
| heading_responsible | string |  |  |

## Data Validation and Quality Control

### Heritabilities of Low-level Traits

Heritability is a statistic used in breeding to quantify how much of a trait’s variation is attributable to the examined genetic material, ranging from 0 to 1. For field phenotyping traits, heritability can indicate the quality of a trait, as one is interested in methods that extract highly genotype-specific values, i.e., traits with high heritability. To assess the quality of low-level traits, heritability was calculated for each time point by setting the genotype factor *g*_*i*_ in Equation 1 to random and estimating genetic and non-genetic variances, as described by Oakey et al. [51]. The resulting heritabilities (Figure 4) follow an expected temporal pattern: increasing with growth and decreasing towards the end of the growth phase. These findings align with previously reported values for the same data set, e.g. 0.77 for senescence traits (2016-2018) [16] and 0.61/0.59 for derived plant height traits (start/stop growth) [12]. Additionally, plant height measurements obtained using TLS have been shown to strongly correlate with drone-based height estimations (correlation of 0.99) [12].

**Figure 4.**
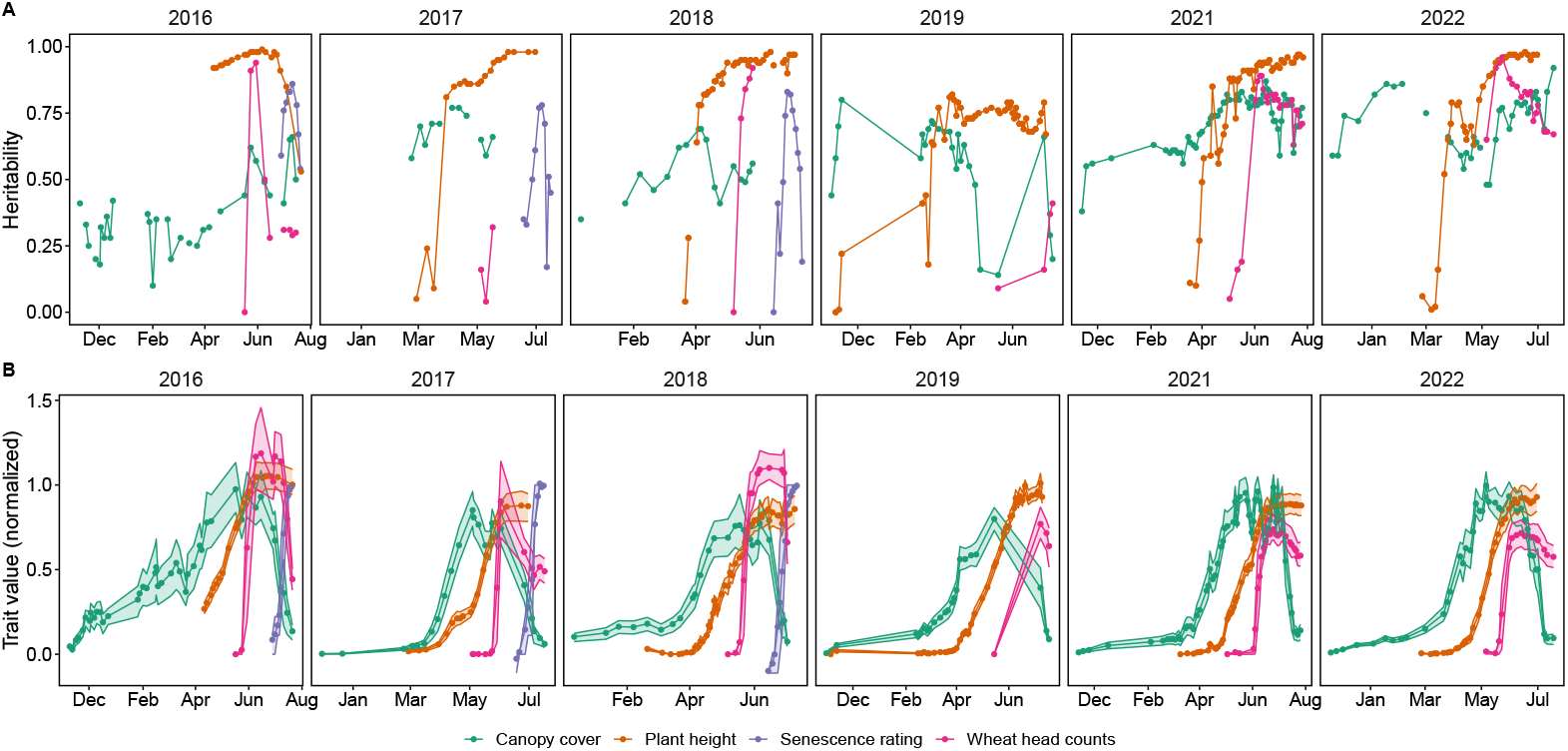
Heritability of the four low-level traits per point in time for all years (2016–2022) (A) and normalized measured trait values (B). Indicated are means (points) and the 25% and 75% percentiles (areas).

**Figure 5.**
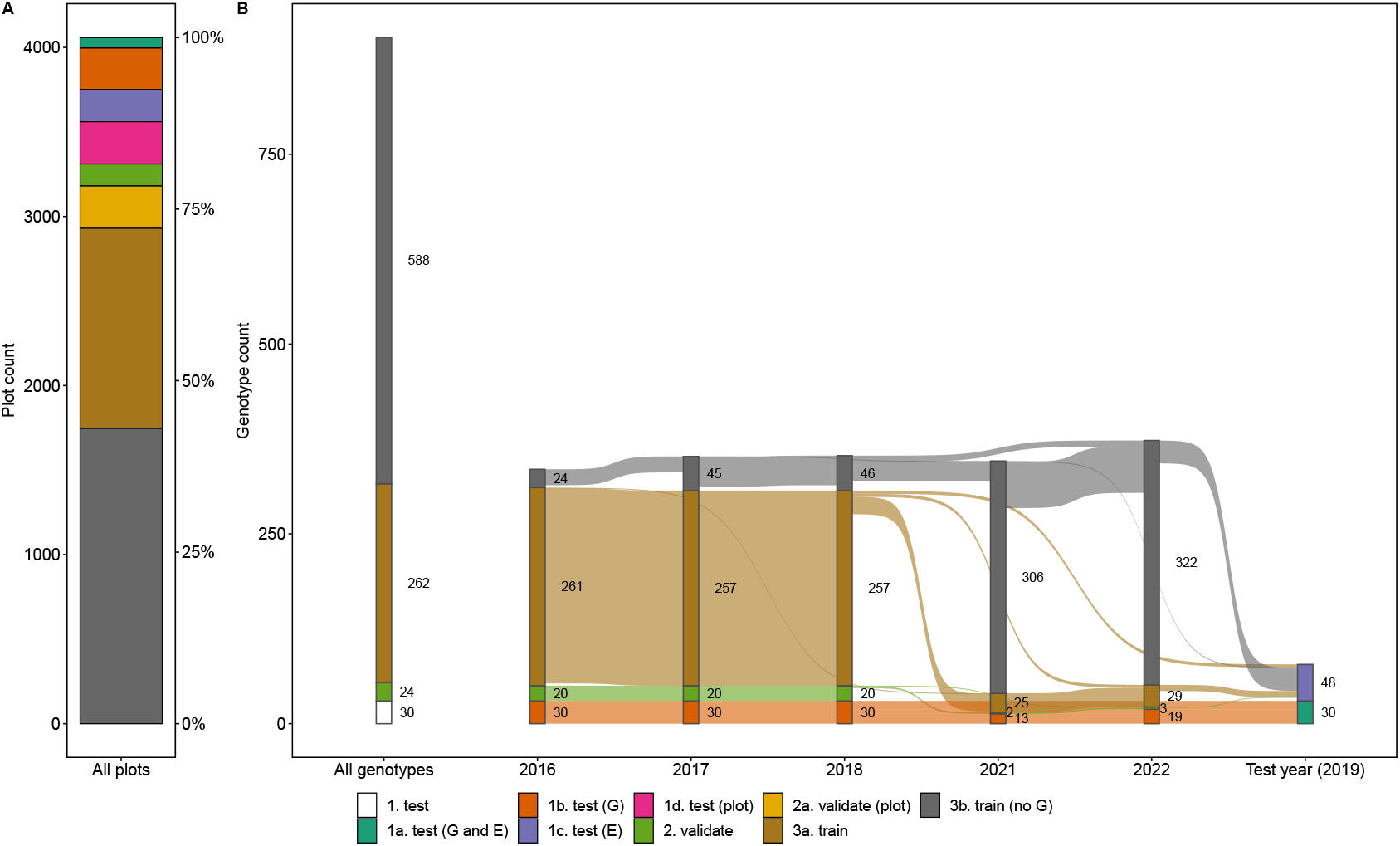
Training, validation and test splits for the GABI-WHEAT marker data based genotype set. The test set includes four subsets, unseen genotypes (G), unseen environments (E), unseen genotypes in unseen environments (G and E), and seen genotypes in seen years on unseen plots (plots). The validation set includes two subsets, unseen genotypes (default) and seen genotypes on unseen plots (plots). The splits are shown based on plots (A) and based on genotype count (B), genotypes occurring in different sets are reported at the lowest level (train < validation < test).

### Heritabilities of Intermediate and Target Traits

To test for the quality of intermediate and target traits, heritability was calculated according to Oakey et al.[51] by setting the genotype factor *g*_*i*_ in Equation 1 to random and estimating genetic and nongenetic variances. The results (Table 5) are in accordance with values reported for the same data set before, e.g., 0.55 for grain yield in 2016–2018 [16], 0.97 for heading date in 2016–2018 [16], 0.84 for grain protein content in 2016 and 2017 [16], and 0.98 for final height in 2015–2018 [12]. The quality of the public marker data set for the GABI-WHEAT panel was demonstrated in Gogna et al. by means of testing for genomic prediction ability [13]. The same quality check was performed on the data set presented herein, once with the public marker data from the GABI-WHEAT panel, once with the extended set that includes private marker data as well.

Three different GBLUP-based genomic prediction models were trained: One linear mixed model with simple main effect and identity variance (ID), one with simple main effect and diagonal variance (DIAG), and one with a simple main effect and random regression to environmental covariates [52]. As environmental covariates, metrics were based on the Standardized Precipitation and Evapotranspiration Index (SPEI) [53], Vapour Pressure Deficit (VPD), air temperature at 2.0 m above ground, and precipitation. For the SPEI and temperature, the mean, maximum and minimum value over the season were used, for VPD, the mean and maximum, and for precipitation, the sum of the values was used. All models were implemented in ASReml-R using code by [54].

Results are reported for the three scenarios unseen genotypes in unseen environment (Test (G and E), Table 9), unseen environments (Test (E), Table 10), and unseen genotypes (Test (G), Table 11). The test (G) corresponds to the scenario reported in Gogna et al. [13]. Results show comparable performance for heading date and final height, superior performance for protein content, and inferior performance for grain yield (Table 11). These results are in accordance to the heritabilities found for the traits (Table 5). If comparing models, no notable difference in performance was found.

**Table 9.** Genomic prediction accuracy (correlation) and bias (RMSE) of intermediate and target traits for unseen genotypes in unseen environments (Test (G and E)).

| Split set | Trait | ID |  |  | DIAG |  |  | RREG |  |  | ID |  |  | DIAG |  |  | RREG |  |  |
| --- | --- | --- | --- | --- | --- | --- | --- | --- | --- | --- | --- | --- | --- | --- | --- | --- | --- | --- | --- |
|  |  | Correlation |  |  |  |  |  |  |  |  | RMSE |  |  |  |  |  |  |  |  |
|  |  | B | CV | MET | B | CV | MET | B | CV | MET | B | CV | MET | B | CV | MET | B | CV | MET |
| GABI | Grain Yield | 0.19 | 0.35 | 0.22 | 0.18 | 0.34 | 0.24 | 0.16 | 0.33 | 0.34 | 1.25 | 1.32 | 1.45 | 1.25 | 1.33 | 1.45 | 1.52 | 1.67 | 1.09 |
| GABI | Grain Yield (adjusted) | 0.16 | 0.34 | 0.21 | 0.16 | 0.34 | 0.23 | 0.15 | 0.33 | 0.31 | 1.57 | 1.67 | 2.52 | 1.57 | 1.67 | 2.51 | 1.86 | 1.99 | 3.1 |
| GABI | Protein Content | 0.36 | 0.46 | 0.46 | 0.34 | 0.45 | 0.46 | 0.33 | 0.43 | 0.46 | 1.11 | 1.13 | 1.23 | 1.13 | 1.14 | 1.23 | 0.65 | 0.67 | 1.55 |
| GABI | Heading Date | 0.59 | 0.55 | 0.67 | 0.59 | 0.55 | 0.67 | 0.59 | 0.55 | 0.67 | 6.57 | 7.31 | 5.77 | 6.54 | 7.28 | 5.77 | 5.35 | 6.08 | 5.86 |
| GABI | Plant Height | 0.8 | 0.75 | 0.81 | 0.8 | 0.74 | 0.81 | 0.57 | 0.61 | 0.75 | 0.06 | 0.07 | 0.08 | 0.06 | 0.06 | 0.08 | 0.18 | 0.17 | 0.17 |
| Extended | Grain Yield | 0.17 | 0.32 | 0.27 | 0.17 | 0.32 | 0.28 | 0.17 | 0.31 | 0.41 | 1.27 | 1.27 | 1.42 | 1.26 | 1.27 | 1.42 | 1.51 | 1.59 | 1.1 |
| Extended | Grain Yield (adjusted) | 0.23 | 0.31 | 0.27 | 0.23 | 0.31 | 0.27 | 0.23 | 0.3 | 0.37 | 1.66 | 1.86 | 2.51 | 1.66 | 1.86 | 2.5 | 1.92 | 2.13 | 3.03 |
| Extended | Protein Content | 0.37 | 0.64 | 0.52 | 0.37 | 0.64 | 0.53 | 0.36 | 0.63 | 0.52 | 1.12 | 1.06 | 1.25 | 1.12 | 1.07 | 1.25 | 0.81 | 0.79 | 1.55 |
| Extended | Heading Date | 0.6 | 0.53 | 0.66 | 0.61 | 0.57 | 0.68 | 0.62 | 0.55 | 0.64 | 6.95 | 6.65 | 5.73 | 6.95 | 6.66 | 5.71 | 5.59 | 5.3 | 5.88 |
| Extended | Plant Height | 0.8 | 0.76 | 0.82 | 0.8 | 0.78 | 0.81 | N/A | 0.65 | 0.8 | 0.05 | 0.05 | 0.08 | 0.05 | 0.05 | 0.08 | N/A | 0.34 | 0.17 |

ID: Identical variances; DIAG: Varying variances per year; RREG: Random regression to environmental covariates; N/A: Failed convergence
Test set: B: Balanced FIP 1.0 data set; CV: 5-fold cross-validation on FIP 1.0 data set; MET: Unseen multi-environment trial data set

**Table 10.**
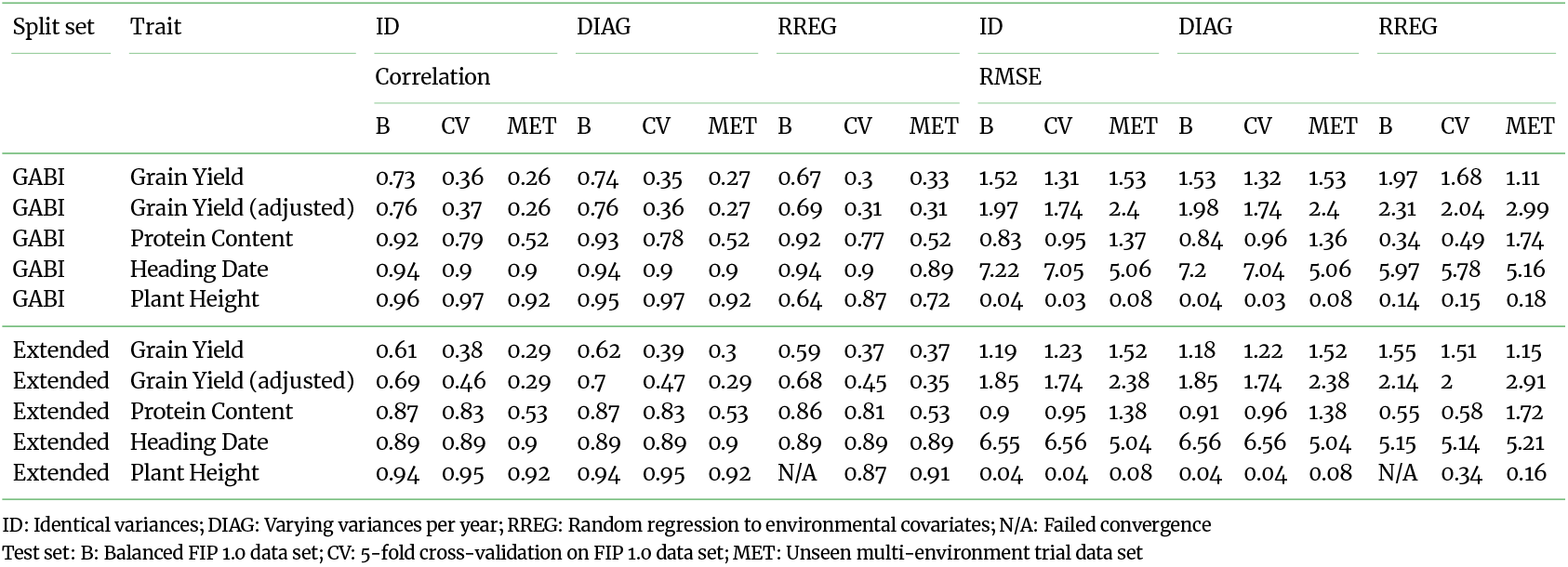
Genomic prediction accuracy (correlation) and bias (RMSE) of intermediate and target traits for seen genotypes in unseen environments (Test (E)).

**Table 11.** Genomic prediction accuracy (correlation) and bias (RMSE) of intermediate and target traits for unseen genotypes in seen environments (Test (G)).

| Split set | Trait | ID |  | DIAG |  | RREG |  | ID |  | DIAG |  | RREG |  |
| --- | --- | --- | --- | --- | --- | --- | --- | --- | --- | --- | --- | --- | --- |
|  |  | Correlation |  |  |  |  |  | RMSE |  |  |  |  |  |
|  |  | B | CV | B | CV | B | CV | B | CV | B | CV | B | CV |
| GABI | Grain Yield | 0.45 | 0.44 | 0.44 | 0.42 | 0.45 | 0.44 | 1.04 | 1 | 1.04 | 1 | 1.04 | 1 |
| GABI | Grain Yield (adjusted) | 0.45 | 0.44 | 0.44 | 0.42 | 0.45 | 0.44 | 0.69 | 0.66 | 0.69 | 0.66 | 0.69 | 0.66 |
| GABI | Protein Content | 0.61 | 0.58 | 0.62 | 0.58 | 0.61 | 0.59 | 0.75 | 0.88 | 0.73 | 0.87 | 0.75 | 0.87 |
| GABI | Heading Date | 0.59 | 0.53 | 0.59 | 0.53 | 0.59 | 0.53 | 2.08 | 2.24 | 2.1 | 2.24 | 2.08 | 2.24 |
| GABI | Plant Height | 0.83 | 0.7 | 0.83 | 0.71 | 0.76 | 0.69 | 0.07 | 0.08 | 0.07 | 0.08 | 0.08 | 0.08 |
| Extended | Grain Yield | 0.48 | 0.37 | 0.47 | 0.39 | 0.48 | 0.37 | 0.99 | 1.04 | 0.99 | 1.04 | 0.99 | 1.04 |
| Extended | Grain Yield (adjusted) | 0.48 | 0.37 | 0.47 | 0.39 | 0.48 | 0.37 | 0.65 | 0.69 | 0.65 | 0.69 | 0.65 | 0.69 |
| Extended | Protein Content | 0.7 | 0.79 | 0.71 | 0.79 | 0.7 | 0.79 | 0.68 | 0.69 | 0.68 | 0.7 | 0.68 | 0.69 |
| Extended | Heading Date | 0.63 | 0.53 | 0.65 | 0.58 | 0.64 | 0.53 | 1.93 | 2.13 | 1.88 | 2.05 | 1.91 | 2.11 |
| Extended | Plant Height | 0.82 | 0.67 | 0.83 | 0.71 | N/A | 0.63 | 0.06 | 0.07 | 0.06 | 0.07 | N/A | 0.07 |

ID: Identical variances; DIAG: Varying variances per year; RREG: Random regression to environmental covariates; N/A: Failed convergence
Test set: B: Balanced FIP 1.0 data set; CV: 5-fold cross-validation on FIP 1.0 data set

### Genomic Prediction Ability of Unseen Multi-environment Trial

Gogna et al. have published a MET data set for yield, protein content, heading date and final height comprising eight environments (five locations and one to two years) for the GABI-WHEAT panel [13]. 312 of the measured genotypes overlap with the FIP 1.0 data set, 60 are unseen in the FIP 1.0 data set but marker data are available. The unseen genotypes (60) in unseen environments (Test (G and E)) and seen genotypes (312) in unseen environments (Test (E)) were taken as independent test sets in new environments for a genomic prediction approach similar to the one described in the previous section. For the random regression model, hourly temperature, precipitation and relative humidity data for the German environments were extracted from the Climate Data Center (CDC) of the German Weather Service, those for the French environments from Meteo France (SYNOP, 3-hourly data only). These additional environmental covariate data and MET data are available in the data repository for convenience, but not part of the core data set (see folder ‘MET_repository_clone’).

For grain yield, the random regression model outperformed the other models (Table 9). For all other traits, no clear advantage of the random regression model over the other models was visible. Again, the results suggest comparable accuracies to Gogna et al. [13] for final height, heading date and protein content, and inferior performance for grain yield (Table 9).

### Re-use Potential and Limitations

In this work, we provide baselines for genomic prediction approaches, trait extractions from images, and subsequent trait dynamics modeling. Accordingly, we see the largest re-use potential of the presented data set for the development and evaluation of new modelling and prediction approaches in crop genomics and phenomics. The multi-faceted data set allows modelling approaches on various levels:

- Genomic prediction approaches that include genotype-environment interactions: The presented data enhance the data by Gogna et al. [13] by 6 environments, totalling to 14 environ-ments that are characterized by environmental covariates. The presented benchmark of a genomic prediction with random regressions to environmental covariates [52] provides a baseline that novel approaches can challenge.
- Modelling plant growth and development with longitudinal modelling approaches: The four low-level traits canopy cover, plant height, wheat head count and senescence cover the full growing season of winter wheat in 6 environments that are characterized by environmental covariates. Baseline approaches for plant height growth modelling [8, 9, 19, 20, 21, 12], canopy cover growth modelling [25] and senescence dynamics modelling [15, 16, 17] for subsets of the presented data exist.
- Image-based phenomic predictions and combined phenomic and genomic prediction approaches: The dense time series of images allow training and analysing end-to-end modelling approaches (e.g., deep learning based) that predict target traits such as yield based on images.

While the data set opens up the possibility of analysing HTFP data to a wide audience, it also has its inherent limitations that should be taken into account if working with it:

- The immobility of the FIP restricts the data set to only one location.
- Field-based data collection introduces various sources of errors that one must consider in analysis (see e.g. [20] for a discussion).
- Yield measurements in the FIP and hence this data set are more prone to error than in METs.
- Annotation at the image level only requires further annotation effort if semantic segmentation or object detection methods are targeted.
- While the aligned image time series provide extensive opportunities to analyze growth dynamics, this kind of highly preprocessed image data is rare and therefore interoperability with other data sources is yet limited.

## Examples

To run the following examples the Huggingface *datasets* [27] library is required. The examples were run using version *3.3.2*.

### Example 1: Basic access to data set via hugging face datasets package

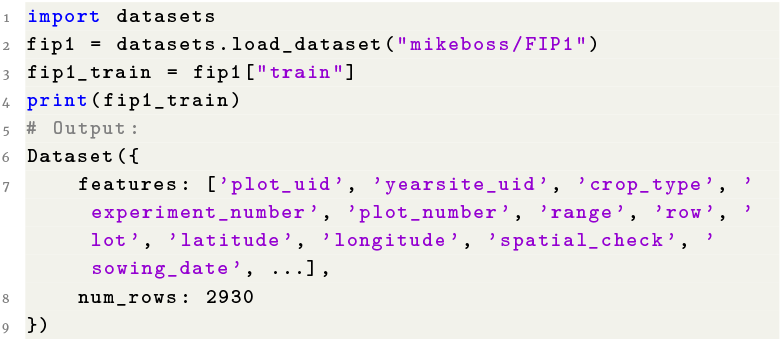

**Listing 1**. Example 1

### Example 2: Load aligned inner plot cutouts as stacked numpy array

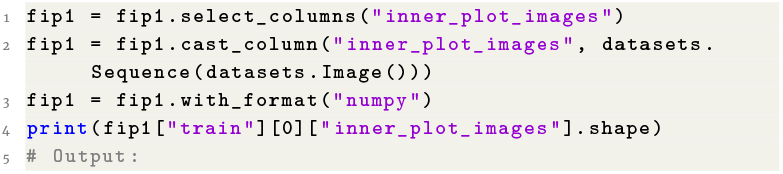
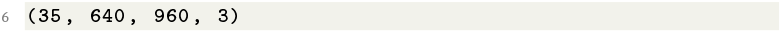

**Listing 2**. Example 2

### Example 3: Access low-level trait time series

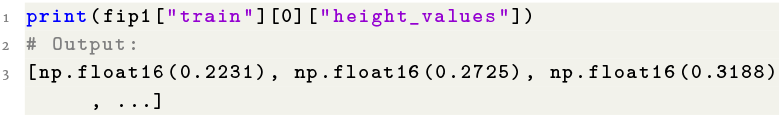

**Listing 3**. Example 3

### Example 4: Access specific target traits

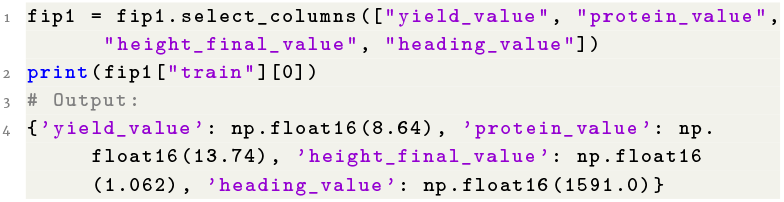

**Listing 4**. Example 4

### Example 5: Access marker data

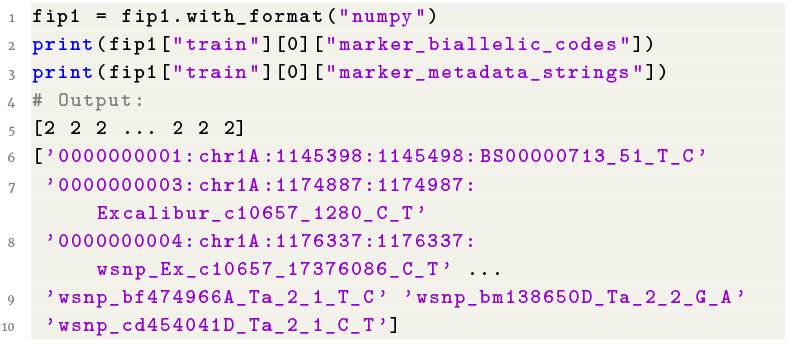

**Listing 5**. Example 5

### Example 6: Access environmental data in January

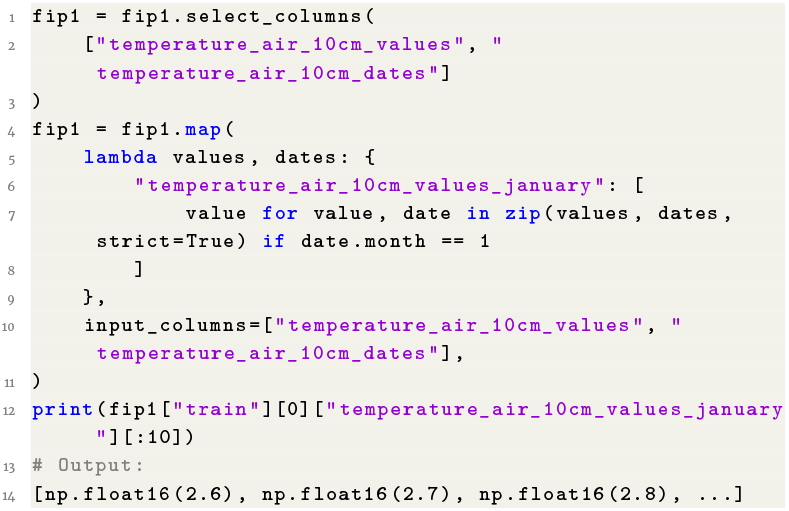

**Listing 6**. Example 6

## Availability of Source Code and Requirements

The code to recreate the derived data and the data set is publicly available in three repositories, namely the *FIP 1.0 Data Set - Traits, fip1-alignment*, and *fip1-dataset* repositories.

The complete process to create the data set involves extracting trait data from the raw data using the *FIP 1.0 Data Set - Traits* repository, then aligning the image time-series using the *fip1-alignment* repository, and finally aggregating the derived data into the final data set using the *fip1-dataset* repository.

In addition, the data set can be recreated using the *fip1-dataset* repository from the derived data that is freely available in the ETH research collection.

### Trait Data Compilation

Project name: FIP 1.0 Data Set - Traits

Project home page: https://gitlab.ethz.ch/crop_phenotyping/fip-1.0-data-set-traits

Operating system(s): Platform independent Programming language: R, Python License: GNU GPL v3

### Image Data Alignment

Project name: fip1-alignment

Project home page: https://gitlab.ethz.ch/crop_phenotyping/fip1-alignment

Operating system(s): Platform independent Programming language: Python

License: GNU GPL v3

### Data Set Compilation

Project name: fip1-dataset

Project home page: https://gitlab.ethz.ch/crop_phenotyping/fip1-dataset

Operating system(s): Platform independent Programming language: Python

License: GNU GPL v3

## Data Availability

- Data Repository: http://doi.org/20.500.11850/697773
- Hugging Face Data set: https://huggingface.co/datasets/mikeboss/FIP1
- Public GABI marker data repository (also integrated in main Data Repository and Hugging Face Data set): https://doi.org/10.5061/dryad.n02v6wwzc
- Private Agroscope marker data repository: Confidential (Contact: Boulos Chalhoub). This repository contains marker data (Illumina Infinium 25k array) from eight generation (F8) breeding lines that are unregistered and property of Agroscope. Access can be requested by stating the intended purpose of use and the willingness to sign a material transfer agreement (MTA).

## Declarations

## Glossary

F8: Eighth Generation Breeding Material. 2, 3, 5
RGB: Red, Green and Blue. 2, 4

## Acronyms

BLUE: Best Linear Unbiased Estimate. 4
CNNs: Convolutional Neural Networks. 2, 5
FIP: Field Phenotyping Platform. 2–4, 6, 9
FIP 1.0: Field Phenotyping Platform 1.0. 2, 3, 6, 14
GABI-WHEAT: Genomanalyse im Biologischen System Pflanze - Weizen. 2, 3, 5, 6, 9, 11
HTFP: High-throughput field phenotyping. 2, 6
INVITE: Innovation in Variety Testing. 2
MET: Multi-Environment Trial. 2, 4, 6
QMER: Quarter of Maximum Elongation Rate. 4
SfM: Structure from Motion. 4
SIFT: Scale-invariant feature transform. 3
SpATS: Spatial Analysis of Field Trials with Splines. 5
SPEI: Standardized Precipitation and Evapotranspiration Index. 6
TLS: Terrestrial Laser Scanner. 2, 4, 5
VPD: Vapour Pressure Deficit. 6

## Consent for Publication

Not applicable.

## Author’s Contributions

Lukas Roth: Conceptualization, Methodology, Software, Validation, Formal analysis, Investigation, Data Curation, Writing - Original Draft, Visualization, Supervision, Funding acquisition. Mike Boss: Conceptualization, Methodology, Software, Validation, Formal analysis, Investigation, Data Curation, Writing - Original Draft, Visualization. Norbert Kirchgessner: Conceptualization, Methodology, Software, Validation, Formal analysis, Investigation, Data Curation, Writing - Original Draft, Visualization. Helge Aasen: Investigation, Supervision. Brenda Patricia Aguirre-Cuellar: Investigation. Price Pius Atuah Akiina: Investigation. Jonas Anderegg: Methodology, Data Curation, Investigation. Joaquin Gajardo Castillo: Software, Data Curation. Xiaoran Chen: Investigation. Simon Corrado: Investigation. Krzysztof Cybulski: Software, Data Curation. Beat Keller: Investigation, Supervision. Stefan Göbel Kortstee: Investigation. Lukas Kronenberg: Methodology, Data Curation, Investigation. Frank Liebisch: Investigation, Supervision. Paraskevi Nousi: Investigation. Corina Oppliger: Investigation. Gregor Perich: Investigation. Johannes Pfeifer: Investigation. Kang Yu: Investigation. Nicola Storni: Software, Data Curation, Investigation. Flavian Tschurr: Software, Data Curation, Investigation. Michele Volpi: Investigation, Supervision. Simon Treier: Investigation, Data Curation. Hansueli Zellweger: Investigation. Olivia Zumsteg: Investigation. Andreas Hund: Conceptualization, Methodology, Writing - Review & Editing, Supervision, Project administration. Achim Walter: Conceptualization, Writing - Review & Editing, Supervision, Project administration, Funding acquisition.

## Acknowledgements

Not applicable.

